## Supplementary Information for "Fast volumetric fluorescence lifetime imaging of multicellular systems using single-objective light-sheet microscopy"

Figure S1 – Benchmarking of soSPIM-FLIM on AF488 dye solution

Table S1 – Comparison light-sheet FLIM and confocal.

Table S2 – Acquisition parameters for soSPIM-FLIM acquisitions

Figure S2 – Comparison to confocal in different depths.

Figure S3 – Confocal reference fits for figure 1.

Figure S4 – Effect of pile-up on dynamic range of images recorded with the SPAD array.

Figure S5 – Adapted Pile-up correction for scanning light-sheet.

Figure S6 – Adapted pile-up correction for soSPIM-FLIM with digital scanned light-sheet.

Figure S7 – Pile-up correction for static light-sheet.

Figure S8 – Confocal reference measurement for lifetime unmixing.

Figure S9 – Single cell fits for Flipper-TR Time-Lapse.

Figure S10 – Dark counts and IRF of the SPAD array detector.

Movie S1 – 3D Multiplexing

Movie S2 – 3D Time-lapse Tension Imaging

#### Comparison soSPIM-FLIM and Confocal

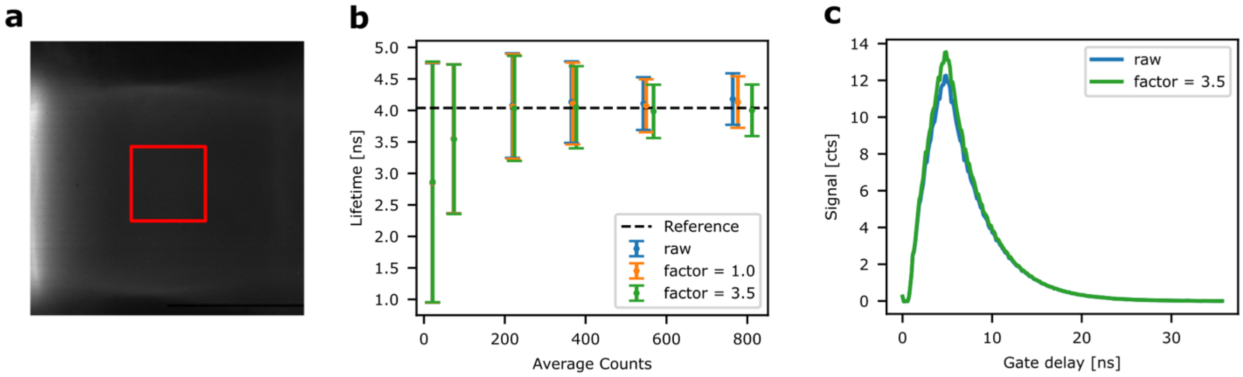


**Figure S1 – Benchmarking of soSPIM-FLIM on AF488 dye solution. (a)** Intensity image of AF488 fluorescence excited using scanning light-sheet, summed over 250 gates, each exposed for 20 ms exposure, shifted with a step size of 143 ps. **(b)** Fluorescence lifetime measured by soSPIM-FLIM with different illumination powers, plotted as a function of average photon count (summed over all gates) in the central 128x128 detector pixels (see red box in (a)). Shown are average lifetimes and standard deviations of all pixels in the central region, analyzed by pixel-wise fitting for three different pile-up corrections (raw: no correction, static correction with factor 1, empirical correction for scanning mode with factor 3.5, see Fig. S5 for details). The dashed black line shows the lifetime of 4.04 ns determined by confocal FLIM performed for 2 min at 2 MHz count rate from fitting the summed TCSPC decay. **(c)** Pixel-averaged decay for the measurement shown in (b) with the highest photon count, before (raw, average lifetime of 4.18 ns) and after pile-up correction with factor 3.5 (average lifetime of 4.00 ns).

**Table S1 – Comparison light-sheet FLIM and confocal.** Power densities, count rates, and extracted lifetimes for the measurements shown in Fig. 1 and Fig. S2. The power densities are calculated for a confocal waist of 0.2 µm (as measured by FCS), waists of 1.46 µm for the scanning Gaussian beam, 1.88 µm waist and 109 µm width for the static light-sheet created by the cylindrical lens. Based on these parameters, which were experimentally determined by imaging the beam and the light-sheet in dye solution, the cross-sectional area of the beam was determined. For details on experiments and analysis see Methods.

| | **Average Laser Power [µW]** | **Power density [kW/cm²]** | **Acqui-sition Time [s]** | **Otsu Thres-hold [cts]** | **Max Counts (99.9%) [cts]** | **Max Pixel Count-rate [kcps]** | **Avg Count-rate [Mcps]** | **Lifetime mean [ns]** | **Lifetime sdev [ns]** |
|---|---|---|---|---|---|---|---|---|---|
| Lightsheet Static depth 30 µm | 770 | 0.2 | 0.10 | 258 | 844 | 8.1 | 414.6 | 2.99 | 0.26 |
| | 770 | 0.2 | 0.18 | 593 | 1946 | 10.8 | 547.6 | 2.96 | 0.18 |
| | 770 | 0.2 | 0.33 | 1206 | 3967 | 12.0 | 606.4 | 2.88 | 0.13 |
| Lightsheet Scanning depth 30 µm | 70 | 1.0 | 0.63 | 127 | 423 | 0.7 | 37.2 | 3.09 | 0.46 |
| | 70 | 1.0 | 1.13 | 255 | 835 | 0.7 | 41.5 | 3.06 | 0.33 |
| Confocal RapidFLIM depth 50 µm | 83 | 66.1 | 1.37 | 6 | 83 | 31,923 | 0.6 | 3.00 | 1.00 |
| | 83 | 66.1 | 10.93 | 45 | 652 | 31,332 | 0.6 | 3.00 | 0.37 |
| | 83 | 66.1 | 34.15 | 139 | 2023 | 31,119 | 0.6 | 3.01 | 0.24 |
| Confocal RapidFLIM depth 20 µm | 83 | 66.1 | 1.37 | 11 | 116 | 44,615 | 0.7 | 2.91 | 0.67 |
| | 83 | 66.1 | 10.93 | 89 | 912 | 43,846 | 0.7 | 2.93 | 0.28 |
| | 83 | 66.1 | 34.15 | 276 | 2808 | 43,200 | 0.7 | 2.93 | 0.21 |
| Confocal standard depth 50 µm | 8.5 | 6.8 | 22.79 | 1 | 16 | 342 | 0.009 | 3.16 | 2.03 |
| | 8.5 | 6.8 | 113.94 | 6 | 73 | 156 | 0.010 | 3.17 | 1.03 |
| Confocal standard depth 20 µm | 8.5 | 6.8 | 22.79 | 3 | 35 | 841 | 0.014 | 3.07 | 1.32 |
| | 8.5 | 6.8 | 113.94 | 16 | 171 | 365 | 0.015 | 3.09 | 0.64 |

**Table S2 – Acquisition parameters soSPIM-FLIM for all measurements.** Acquisition time is per FLIM frame, acquiring all gate delay steps including overhead for 3D acquisitions (Multiplexing and Flipper TR).

| | **Acquisition Time [s]** | **Gate delay steps** | **Delay Step [ns]** | **Exposure per delay step [ms]** | **Average Laser Power [µW]** |
|---|---|---|---|---|---|
| E-Cadherin-AF488 (static) | 0.10 | 25 | 0.609 | 3 | 770 |
| | 0.18 | 50 | 0.304 | 3 | 770 |
| | 0.33 | 100 | 0.143 | 3 | 770 |
| E-Cadherin-AF488 (scanning) | 0.63 | 25 | 0.501 | 20 | 70 |
| | 1.13 | 50 | 0.251 | 20 | 70 |
| Multiplexing GFP / Flipper-TR | 1.13 | 50 | 0.358 | 20 | 70 |
| | 2.32 | 100 | 0.179 | 20 | 70 |
| Flipper Org 1 | 1.13 | 50 | 0.358 | 20 | 70 |
| Flipper Org 2 | 2.83 | 50 | 0.358 | 50 | 30 |
| Flipper Org 3 | 1.13 | 50 | 0.358 | 20 | 120 |

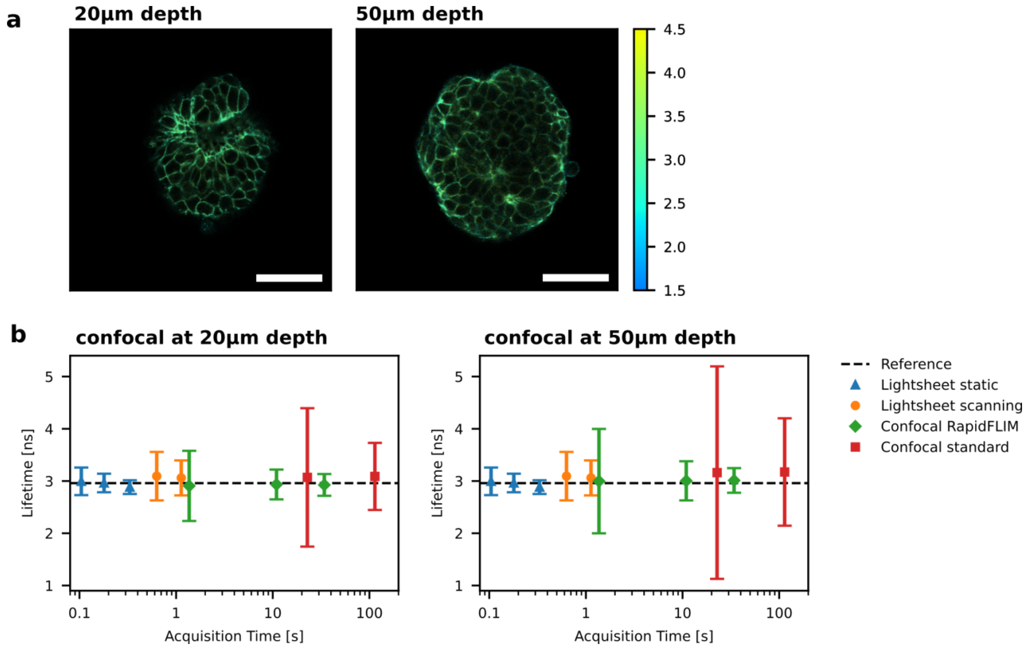


**Figure S2 – Comparison to confocal in different depths. (a)** Example confocal images of fixed embryonic organoids stained with E-cadherin-AF488 at different depths, showing the loss in intensity of around 25-30% at higher depth inside the organoid (see also Fig. S3). Scale bars 50µm. **(b)** Lifetime accuracy of light-sheet FLIM at 30 µm depth compared to confocal measurements at 20 µm and 50 µm depth. In line with the higher count rate, the confocal lifetime histograms are narrower at lower depth.

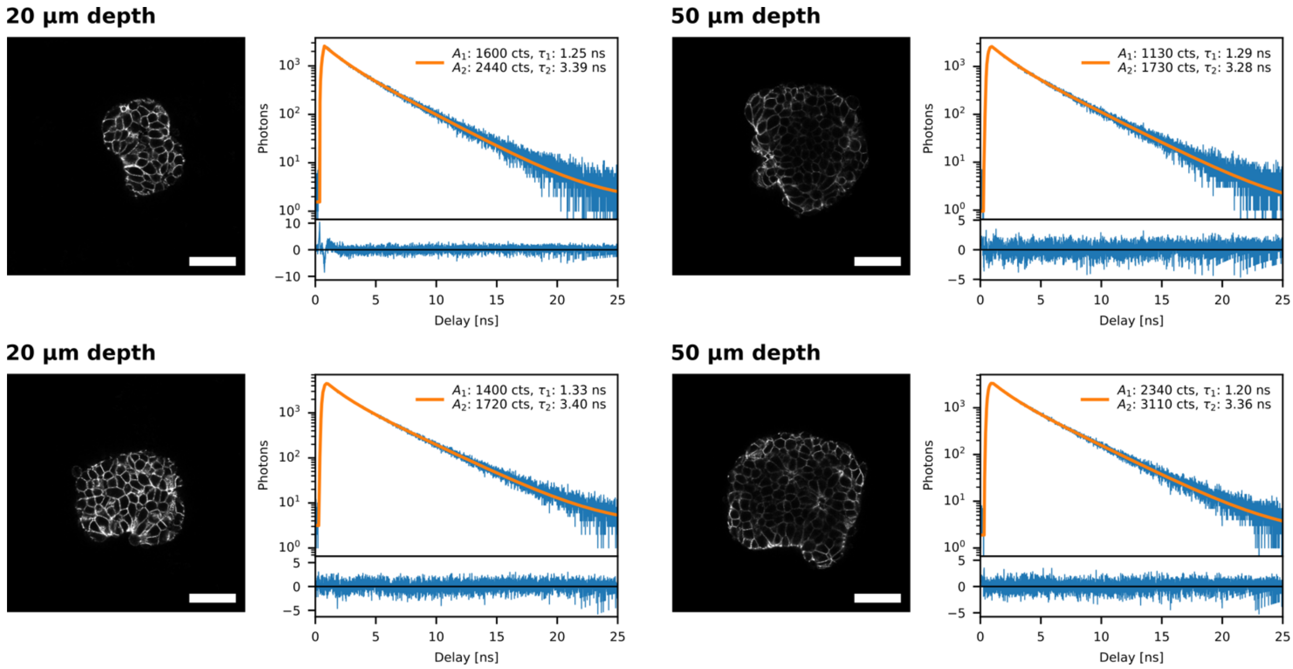


**Figure S3 – Confocal reference fits for figure 1.** Intensity images (scale bars 50 µm) and sum fits of fixed embryonic organoids stained with E-cadherin-AF488 at depths of 20 µm and 50 µm inside the organoid. The intensity weighted average lifetimes from the biexponential fits are used as the reference lifetime for Fig. 1c.

### Pile-up correction and dynamic range

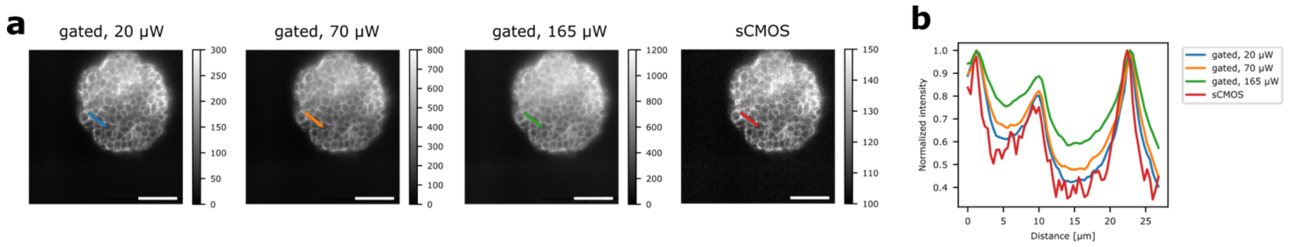


**Figure S4 – Effect of pile-up on dynamic range of images recorded with the SPAD array. (a)** Intensity images of Ecad-AF488, calculated by summing the photon counts over all 50 time gates with 20ms exposure per gate, acquired at different laser powers using scanning light-sheet. The rightmost image was recorded with an sCMOS camera, using 70 µW excitation power and 20 ms total exposure. (scale bars 50 µm) **(b)** Normalized intensity profile across the lines drawn in (a). At low laser power the profiles from SPAD array and sCMOS images are similar, the latter showing more noise due to shorter total exposure. At higher laser power, the SPAD array saturates due to limited dynamic range.

Due to the working principle of the time-gated SPAD array detector, only 1 photon can be detected per pixel within a single 1-bit exposure (minimum exposure time ca. 10 µs). Additional photons incident on the same pixel during the exposure period are not detected. This leads to pile-up effects in bright image regions and the loss of dynamic range. For 8-bit images (summed over 255 1-bit exposures) this can be compensated with the formula

$$N_{corr} = -\ln(1 - N/255) * 255,$$

if it is not to severe. However, for data acquisition in the digital scanned light-sheet modality, each pixel is only illuminated for a fraction of the overall exposure and fewer 1-bit exposures can contribute to meaningful signal. Therefore, the usable dynamic range of the sensor is limited more severely and saturation sets in before the detector reaches its nominal saturation count rate (Fig. S4). Additionally, achievable imaging speed is limited by the scan rate, which was 1 kHz over the field of view (monodirectional) in this case. The typically used acquisition settings were 8-bit exposure times of 20 ms (i.e. 20 monodirectional scans over the field of view) and maximum count rates of 40 counts per pixel in an 8-bit image. In figure S4 this corresponds to the image recorded with 70 µW average laser power. At these settings, we observe pile-up evident as loss of dynamic range (Fig. S4). To compensate for the reduced number of 1-bit exposures contributing meaningful signal and additional photons in a 1-bit exposure not being detected, we adapted the pile-up correction formula by adding an additional scaling factor F:

$$N_{corr} = -\ln(1 - N * F/255) * 255/F$$

This factor models that a certain photon count in an 8-bit image is generated by only a fraction of 1-bit gate exposures and thus locally higher photon count rates, compared to the conventional pile-up correction given above.

In gated imaging, pile-up will mostly affect the intensity for the gate delays with high count rates, corresponding to short fluorescence lifetimes (Fig. S5). Thus, contrary to classical pile-up in TCSPC measurements, pile-up leads to systematically longer lifetime estimates. We use this correlation between intensity and measured lifetime in scanning light-sheet measurements of ecad-AF488 labelled fixed organoids to determine an empirical correction factor F for our experiments. We analyze FLIM images acquired in scanning mode at different laser powers with a range of correction factors and minimize the above mentioned correlation between count rate and fluorescence lifetime (Fig. S5).

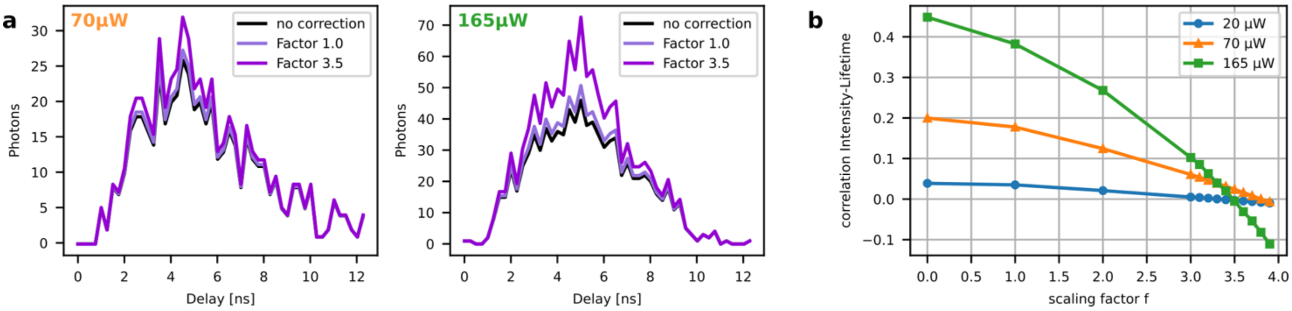


**Figure S5 – Adapted Pile-up correction for scanning light-sheet. (a)** Effect of standard (F=1) and adapted (F=3.5) pile-up correction on typical in-focus single pixel decays at different laser powers. The adapted pile-up correction leads to a total increase in counts of approximately 13 % and 30 % at 70 µW and 165 µW respectively. **(b)** Pearson correlation of pixel intensity and pixel lifetime from mono-exponential fit over the scaling factor F for lifetime images obtained at 3 different laser powers.

We found that a scaling factor of 3.5 can minimize correlation across laser powers (Fig. S5). After the adapted pile-up correction, the correlation between intensity and lifetime in scanning light-sheet FLIM images is mostly removed (Fig. S6). We found that the scaling factor is largely independent of laser power and depends predominantly on scanning settings, in particular the scanning frequency and the effective width of the scanning beam and thereby generated fluorescence. However, the latter is not uniform across the sample (within the image plane and at different depth in the sample), not only due to the Gaussian nature of the beam, but in particular due to aberrations and scattering in optically challenging samples such as the organoids measured here. This effectively restricts the spatial region where the empirical pile-up correction is valid. Global optimization across the whole image plane as performed here thus may still suffer from local pile-up in the region of the beam waist, as seen in Fig. 2.

In contrast to the scanning light-sheet, the full dynamic range and imaging speed of the SPAD-array can be used in FLIM with the static light-sheet configuration. Typical acquisition settings for scanning light-sheet were 3 ms exposure per time-gate and up to 100 counts in the brightest pixels within one 8-bit image. In static light-sheet FLIM the already weak correlation between pixel intensity and lifetime can be compensated by the standard pile-up correction with correction factor F=1 (Fig. S7).

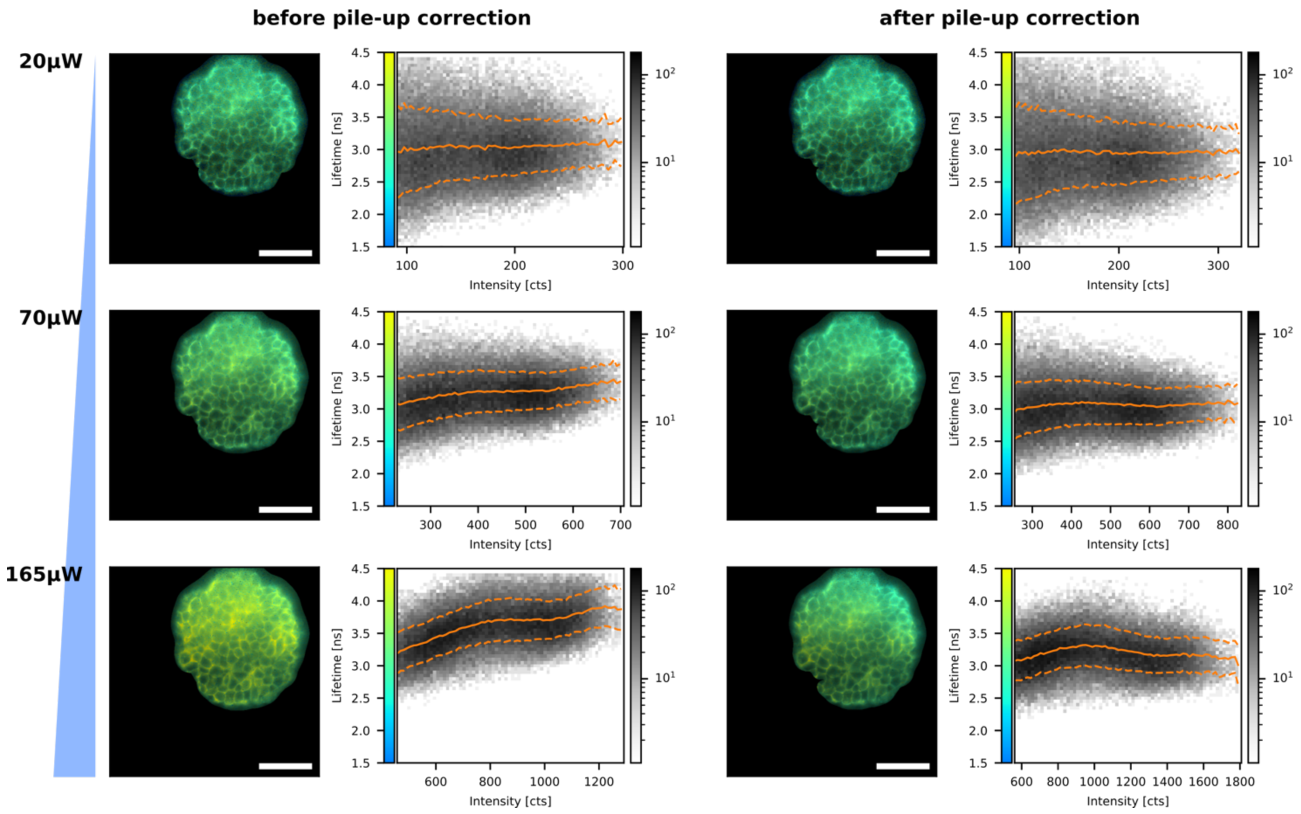


**Figure S6 – Adapted pile-up correction for soSPIM-FLIM with digital scanned light-sheet.** Scanning soSPIM-FLIM images of fixed embryonic organoids stained with E-cadherin-AF488 and 2D histograms (pixel lifetime over pixel intensity) before and after adapted pile-up correction with correction factor F=3.5. The orange lines show the lifetime and standard deviation over the pixel counts.

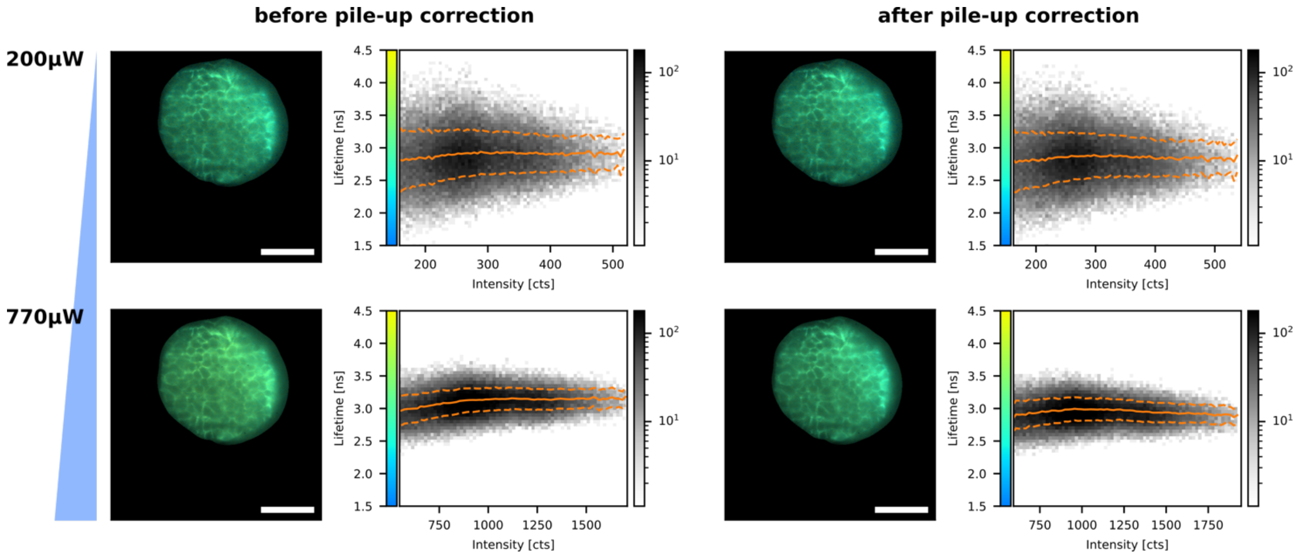


**Figure S7 – Pile-up correction for static light-sheet.** Static soSPIM-FLIM images of fixed embryonic organoids stained with E-cadherin-AF488 and 2D histograms (pixel lifetime over pixel intensity) before and after standard pile-up correction (correction factor F=1).

### Applications

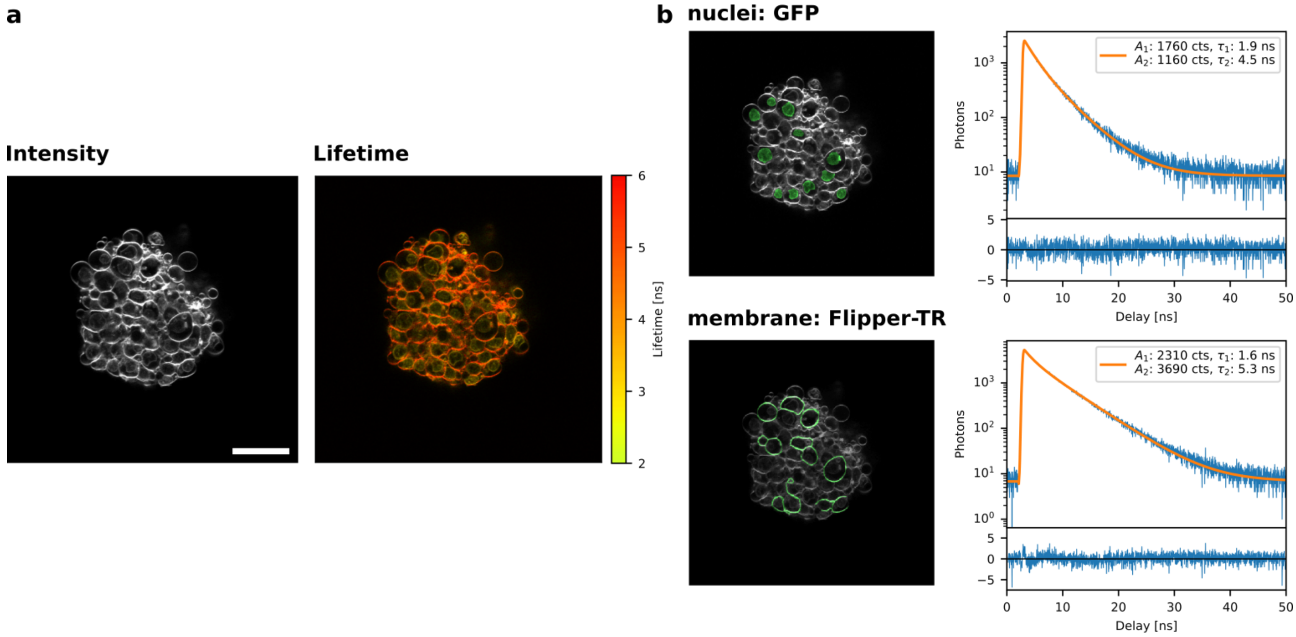


**Figure S8 – Confocal reference measurement for lifetime unmixing. (a)** Exemplary confocal intensity and FLIM images of organoids expressing H2B-GFP and additionally stained with Flipper-TR. Scale bar 50 µm. **(b)** Manual selection of regions of interest for nuclei and membranes resulted in species patterns that are fitted with a bi-exponential decay to obtain reference values for Fig. 3b. The intensity weighted average lifetimes for the 2 species are 3.5 ns and 4.7 ns for the nuclei and membrane respectively.

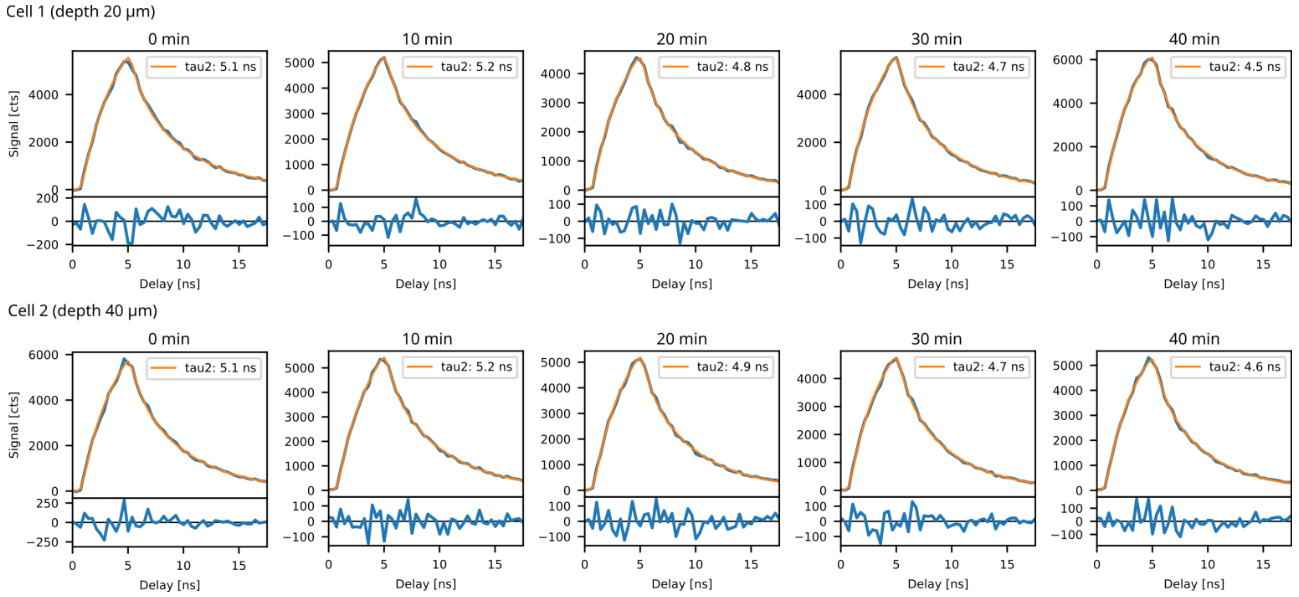


**Figure S9 – Single cell fits for Flipper-TR Time-Lapses.** Shown are example lifetime decays (blue) detected on individual cells (see Fig. 4c) and two-component fits (orange solid lines). Insets show residuals.

### Background and Instrument response function

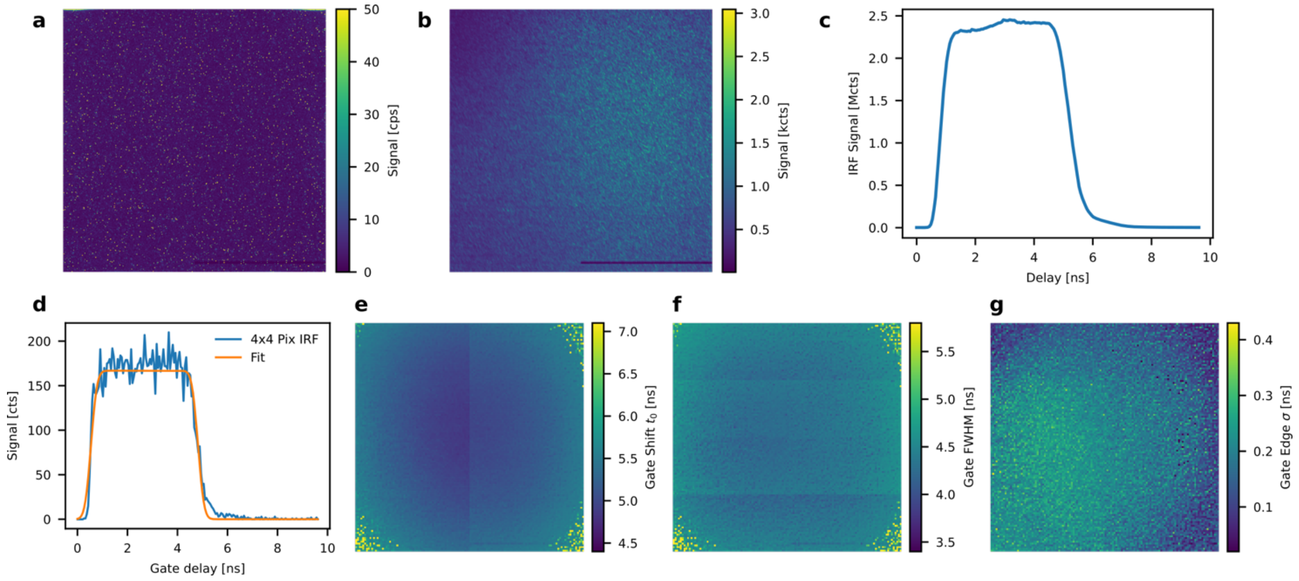


**Figure S10 – Dark counts and IRF of the SPAD-array detector. (a)** Dark counts in gated mode averaged over 180 gate delays and a gate delay step size of 54 ps with 2.7 ms exposure per gate delay. **(b)** IRF measurement intensity summed over all gate steps with the same parameters as in (a). **(c)** Gating signal of the IRF measurement summed over all pixels. **(d)** Example IRF fit binned over 4x4 pixels. **(e)-(g)** Measured IRF parameters across the detector area, namely gate shift of the right flank $t_0$ (e), gate width $w$ (f) and gate edge steepness $\sigma$ (g).

**Movie S1 – 3D Multiplexing.** The video shows soSPIM-FLIM imaging of an organoid in 3D, multiplexed via the lifetime. Membranes are stained with Flipper-TR and nuclei are labelled with GFP. Firstly, intensity images in a few z-planes are shown. In one z-plane, a FLIM-image and the subsequent multiplexed image is shown (see methods). We then show the multiplexed images across the entire depth range. Acquisition time is 1.1 s per 2D plane and the scale bar is 25µm.

**Movie S2 – 3D Lime-lapse Tension Imaging**. The video shows 3D time-lapse FLIM of a live organoid stained with Flipper-TR, first showing 2d planes in increasing depth and then the change of one plane over time. The lifetime scale corresponds to the long lifetime component of a two component decay with the short lifetime fixed (see methods). Scale bar is 25µm.
